# Context-dependent inversion of the response in a single sensory neuron type reverses olfactory preference behavior

**DOI:** 10.1101/2021.11.08.467792

**Authors:** Munzareen Khan, Anna H. Hartmann, Michael P. O’Donnell, Madeline Piccione, Pin-Hao Chao, Noelle D. Dwyer, Cornelia I. Bargmann, Piali Sengupta

**Affiliations:** Department of Biology, Brandeis University, Waltham, MA; Department of Cell Biology, University of Virginia, Charlottesville, VA; The Rockefeller University, New York, NY; Department of Neurobiology, Harvard Medical School, Boston, MA; Department of Molecular, Cellular and Developmental Biology, Yale University, New Haven, CT; Deallus Consulting, New York, NY

## Abstract

The valence and salience of individual odorants are modulated by an animal’s innate preferences, learned associations, and internal state, as well as by the context of odorant presentation. The mechanisms underlying context-dependent flexibility in odor valence are not fully understood. Here we show that the behavioral response of *C. elegans* to bacterially-produced medium-chain alcohols switches from attraction to avoidance when presented in the background of a subset of additional attractive chemicals. This context-dependent reversal of odorant preference is driven by cell-autonomous inversion of the response to alcohols in the single AWC olfactory neuron pair. We find that while medium-chain alcohols inhibit the AWC olfactory neurons to drive attraction, these alcohols instead activate AWC to promote avoidance when presented in the background of a second AWC-sensed odorant. We show that these opposing responses are driven via engagement of different odorant-directed signal transduction pathways within AWC. Our results indicate that context-dependent recruitment of alternative intracellular signaling pathways within a single sensory neuron type conveys opposite hedonic valences, thereby providing a robust mechanism for odorant encoding and discrimination at the periphery.

## INTRODUCTION

Organisms live in complex and dynamic chemical environments. Animals continuously encounter heterogenous mixtures of chemicals which fluctuate in their concentrations and temporal properties, and provide information about the presence and location of food, mates, competitors, and predators (Baker et al., 2018; Laurent, 2002). A particularly critical task of the olfactory system is to discriminate among related chemical cues. Since chemicals that are structurally similar can have distinct saliences for an organism (Bentley, 2006), these cues must be identified and differentiated in order to elicit the appropriate behavior. In particular, animals need to distinguish individual chemicals in a complex mixture, or detect the presence of a new chemical in the background of a continuously present odorant [eg. (Badeke et al., 2016; Livermore and Laing, 1998; Riffell et al., 2014; Rokni et al., 2014)]. Context-dependent odorant discrimination can be driven via integration and processing of multiple sensory inputs in central brain regions [eg. (Groschner and Miesenbock, 2019; Hildebrand and Shepherd, 1997; Kadohisa and Wilson, 2006; Mohamed et al., 2019; Parnas et al., 2013; Stettler and Axel, 2009)]. Although mechanisms operating at the level of single sensory neuron types or sensilla in the periphery have also been implicated in this process (Cao et al., 2017; Duan et al., 2020; Inagaki et al., 2020; Pfister et al., 2020; Reddy et al., 2018; Su et al., 2011; Turner and Ray, 2009; Zhang et al., 2019), the contributions of sensory neurons to mediating odorant discrimination and olfactory behavioral plasticity are not fully understood.

*C. elegans* senses and navigates its complex chemical environment using a small subset of sensory neurons (Perkins et al., 1986; Ward et al., 1975; White et al., 1986). The valence of individual chemicals is largely determined by the responding sensory neuron type, such that distinct subsets of chemosensory neurons drive either attraction or avoidance to different chemicals (Bargmann et al., 1993; Ferkey et al., 2021). Each chemosensory neuron type in *C. elegans* expresses multiple chemoreceptors that are likely tuned to different odorants, a subset of which can be behaviorally discriminated (Troemel et al., 1995; Vidal et al., 2018). Failure to discriminate between two attractive odorants sensed by a single chemosensory neuron results in loss or reduced attraction, but not aversion, to the test chemical (Bargmann et al., 1993). In contrast, aversion of a normally attractive chemical occurs as a consequence of modulation by experience and internal state (Colbert and Bargmann, 1997; Ghosh et al., 2016; Ishihara et al., 2002; Nuttley et al., 2002; Rengarajan et al., 2019; Saeki et al., 2001; Tomioka et al., 2006). Experience- and state-dependent switches in the valence of a chemical occur largely via integration and plasticity at the synaptic and circuit level (Ha et al., 2010; Jang et al., 2012; Kunitomo et al., 2013; Tsunozaki et al., 2008; Yoshida et al., 2012; Zhang et al., 2005), and have not been reported to be driven solely by plasticity in sensory neuron responses.

Here we report that a context-dependent inversion of the odorant response in a single sensory neuron type switches the valence of the behavioral response of *C. elegans* to medium-chain alcohols from attraction to aversion. We show that adult *C. elegans* hermaphrodites are attracted to low concentrations of bacterially produced medium-chain alcohols such as 1-hexanol and 1-heptanol (henceforth referred to as hexanol and heptanol, respectively). However, when presented in the context of a uniform saturating background of a subset of other attractive odorants, animals are now strongly repelled by these alcohols. While the single AWC olfactory neuron pair is known to mediate attraction to a subset of alcohols, we find that in the context of saturating odorants, the AWC neurons instead drive aversion. Hexanol and heptanol inhibit AWC in non-saturating conditions to drive attraction, but in saturating conditions, these chemicals activate AWC to promote avoidance. We show that this context-dependent inversion of response direction is mediated by distinct downstream intracellular effectors including Gα proteins and receptor guanylyl cyclases in AWC. Results described here indicate that contextdependent engagement of distinct intracellular signaling pathways within a single sensory neuron type is not only sufficient to discriminate between structurally related chemicals, but to also convey opposing hedonic valences.

## RESULTS

### Medium-chain alcohols can either attract or repel *C. elegans* based on odorant context

Cross-saturation assays have previously been used as a measure of the ability of *C. elegans* to behaviorally discriminate between two volatile odorants (Bargmann et al., 1993). In these assays, animals are challenged with a point source of the test chemical in the presence of a uniform concentration of a second saturating chemical (Figure 1A). The ability to retain responses to the test chemical under these conditions indicates that the animal is able to discriminate between the test and saturating odorants.

**Figure 1.**
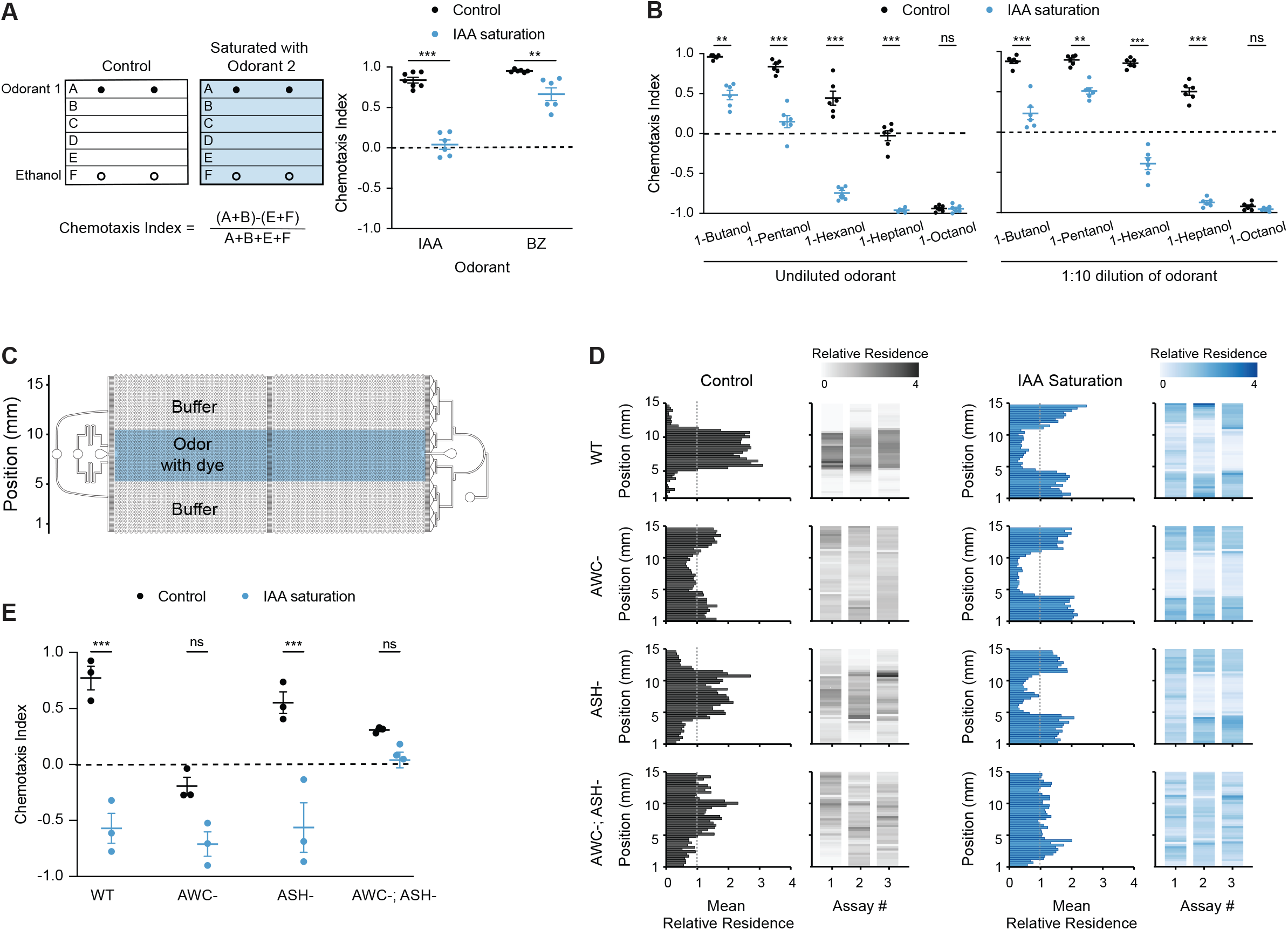
The AWC neurons drive attraction or avoidance of hexanol in the absence or presence of sIAA, respectively. **A)** (Left) Cartoon of plate chemotaxis assay (see Materials and Methods). Filled and open circles: location of 1 μl each of the test odorant or ethanol. Positive and negative chemotaxis indices indicate attraction and avoidance, respectively. (Right) Behaviors of wild-type animals on control or sIAA plates. Test odorants: 1:200 dilution of benzaldehyde (BZ), 1:1000 dilution of IAA. **B)** Behaviors of wild-type animals on control and sIAA plates containing the indicated dilutions of different alcohols as the test odorant. **C)** Schematic of the microfluidics behavioral device used in behavioral assays. Adapted from (Albrecht and Bargmann, 2011). **D)** Average histograms showing mean relative *x-y* residence (relative to spatial odor pattern) of animals of the indicated genotypes over 20 minutes in devices with a central stripe of 10^−4^ hexanol without (left), or with, a uniform concentration of 10^−4^ IAA (right) in the device. Mean relative residence >1 or <1 (dashed vertical line) indicate attraction and avoidance, respectively. Corresponding heat maps show the density of tracks in the *y*-position for each assay. n=20-30 animals per assay; 3 biological replicates. **E)** Chemotaxis indices calculated from behavioral assays shown in **D**. In **A,B**, each dot is the chemotaxis index of a single assay plate containing ~100-200 adult hermaphrodites. Assays were performed in duplicate over at least 3 days. In **E**, each dot is the chemotaxis index from a single assay in behavior chips. Long horizontal bars indicate the mean; errors are SEM. **, ***: *P*<0.01 and 0.001, respectively (**A,B**: Kruskal-Wallis with posthoc pairwise Wilcoxon test and Benjamini-Hochberg method for *P*-value correction; **E**: two-way ANOVA with Bonferroni’s correction); ns: not significant.

The AWC olfactory neuron pair in *C. elegans* responds to low concentrations of a subset of bacterially produced attractive chemicals including benzaldehyde, isoamyl alcohol (IAA), and the short-chain alcohol 1-pentanol (Bargmann et al., 1993; Chalasani et al., 2007; Yoshida et al., 2012). In the presence of a uniform background concentration of IAA, animals are indifferent to a point source of IAA, but retain the ability to respond to a point source of the structurally distinct chemical benzaldehyde (Figure 1A) (Bargmann et al., 1993). To investigate the extent to which AWC is able to discriminate among structurally related chemicals, we examined responses to point sources of straight-chain alcohols with or without saturating IAA (henceforth referred to as sIAA). Attractive responses to the short-chain alcohols 1-butanol and 1-pentanol were reduced or abolished in sIAA indicating that animals are largely unable to discriminate between these chemicals (Figure 1B) (Bargmann et al., 1993). *C. elegans* is robustly repelled by long-chain alcohols such as 1-octanol (Bargmann et al., 1993); this response is mediated by integration of sensory inputs from multiple sensory neurons including AWC in a food-dependent manner (Chao et al., 2004; Summers et al., 2015; Troemel et al., 1995). However, sIAA had no effect on the avoidance of this chemical (Figure 1B). These observations indicate that in sIAA, AWC-driven attractive responses to a subset of related alcohols is decreased or abolished, but that long-range avoidance of alcohols driven by other sensory neurons is unaffected.

Animals were also attracted to the medium-chain alcohols hexanol and heptanol (Figure 1B) (Bargmann et al., 1993). Unlike the reduced attraction observed for short-chain alcohols in sIAA, animals in sIAA strongly avoided both hexanol and heptanol (Figure 1B). In contrast, saturation with either hexanol or heptanol abolished attraction to a point source of IAA but did not result in avoidance (Figure S1A). Many chemicals elicit distinct behaviors at different concentrations [eg. (Bargmann et al., 1993; Horio et al., 2019; Saraiva et al., 2016; Stensmyr et al., 2003)]. Similarly, worms were attracted to low, but were weakly repelled by high, concentrations of hexanol (Figure S1B). The response was shifted towards indifference of lower, and strong avoidance of higher, hexanol concentrations in sIAA (Figure S1B). We infer that distinct underlying antagonistic neuronal pathways mediate attraction to, and avoidance of, hexanol at all concentrations. Attraction predominates at lower, and avoidance at higher, hexanol concentrations; sIAA inhibits the attraction but not the avoidance pathway. These results also indicate that animals are unable to discriminate between IAA and hexanol for attraction but are able to do so for avoidance.

### The attraction-promoting AWC olfactory neuron pair instead drives hexanol avoidance in odorant saturation conditions

The ability of hexanol and IAA to cross-saturate for attraction suggested that hexanol attraction is also mediated by AWC. The AWB, ASH, and ADL sensory neuron pairs mediate avoidance of noxious alcohols including high concentrations of IAA (Ferkey et al., 2021). We tested whether hexanol attraction and avoidance requires AWC and one or more of the avoidance-mediating sensory neurons, respectively.

To more reliably characterize the contributions of different sensory neurons to hexanol attraction and avoidance behaviors, we used microfluidics behavioral arenas (Figure 1C) which enable precise spatiotemporal control of stimulus delivery together with automated tracking of individual worm movement (Albrecht and Bargmann, 2011). Animals were distributed throughout the device in buffer alone, but accumulated within a spatially restricted stripe of IAA or hexanol over the assay period indicating attraction (Figure 1D,E, Figure S1C,D, Video S1). In the presence of a uniform low concentration of IAA throughout the device, animals were indifferent to a central stripe of IAA indicating response saturation (Figure S1C,D). In contrast, in sIAA, animals avoided the central hexanol stripe (Figure 1D,E, Video S1). Thus, the behaviors in microfluidics devices recapitulate the behavioral responses observed in plate chemotaxis assays.

Animals in which AWC was either genetically ablated or silenced via the expression of a gain-of function allele of the *unc-103* potassium channel (Reiner et al., 2006) were no longer robustly attracted to hexanol but instead exhibited weak avoidance, indicating that AWC is necessary for hexanol attraction (Figure 1D,E, Figure S1E). AWC-ablated animals also failed to be attracted to the AWC-sensed odorant benzaldehyde but were not repelled by this chemical (Bargmann et al., 1993) (Figure S1F). In sIAA, AWC-ablated or silenced animals continued to robustly avoid hexanol suggesting that this aversion is mediated by a neuron type other than AWC (Figure 1D,E, Figure S1E). Ablation of the nociceptive neuron type ASH resulted in a failure to avoid high concentrations of the ASH-sensed chemical glycerol (Figure S1G) (Bargmann et al., 1990), but had more minor effects on attraction to, or avoidance of, hexanol without or with sIAA, respectively (Figure 1D,E). However, animals lacking both AWC and ASH were indifferent to hexanol regardless of conditions (Figure 1D,E). We conclude that while AWC drives attraction to hexanol, either AWC or ASH can mediate hexanol avoidance in sIAA. Thus, the typically attraction-mediating AWC sensory neuron pair is able to drive hexanol avoidance based on odorant context.

### Saturation with AWC-sensed odorants cell-autonomously inverts the hexanol response in AWC

To investigate the mechanisms by which AWC mediates hexanol attraction or avoidance in a context-dependent manner, we next examined hexanol-evoked changes in intracellular calcium dynamics. Attractive odorants decrease intracellular calcium levels in AWC (Chalasani et al., 2007). Odorant mediated inhibition of AWC as well as disinhibition and/or activation upon odorant removal together drive attraction (Chalasani et al., 2007; Gordus et al., 2015). We first confirmed that the transgenic strain expressing the genetically encoded calcium indicator GCaMP3 in AWC exhibited behavioral responses to hexanol similar to those of wild-type animals (Figure S2A). We next assessed AWC calcium dynamics in animals immobilized in microfluidics imaging chips (Chronis et al., 2007) in response to hexanol and sIAA concentrations that elicited attraction and avoidance behaviors in microfluidics behavioral arenas described above.

As reported previously (Chalasani et al., 2007), low IAA concentrations that robustly attracts wild-type animals decreased, and removal increased, intracellular calcium levels in AWC (Figure S2B). In sIAA, an additional pulse of IAA led to only a further minor decrease in calcium levels in AWC upon odor onset (Figure S2B), correlating with loss of attraction to IAA in these saturation conditions. Addition of low hexanol concentrations that attracts wild-type animals also decreased calcium levels in AWC (Figure 2A,B, Video S2). However, in sIAA, hexanol instead robustly increased intracellular calcium in AWC correlating with hexanol avoidance under these conditions (Figure 2A,B, Video S2). Animals also avoid heptanol in sIAA (Figure 1B); the heptanol response in AWC was also inverted in this context (Figure 2C). The AWC neuron pair exhibits bilateral response asymmetry to a subset of odorants (Bauer Huang et al., 2007; Chalasani et al., 2007; Wes and Bargmann, 2001). ~90% of imaged AWC neurons responded to both hexanol and heptanol with and without sIAA (Figure 2A,C), implying that both AWC neurons are able to respond to these chemicals regardless of conditions. Hexanol-driven calcium increases and decreases in AWC were maintained in *unc-13* and *unc-31* mutants that lack synaptic and peptidergic transmission, respectively (Sieburth et al., 2007; Speese et al., 2007) (Figure 2B, Figure S2C), suggesting that these responses are mediated cell-autonomously.

**Figure 2.**
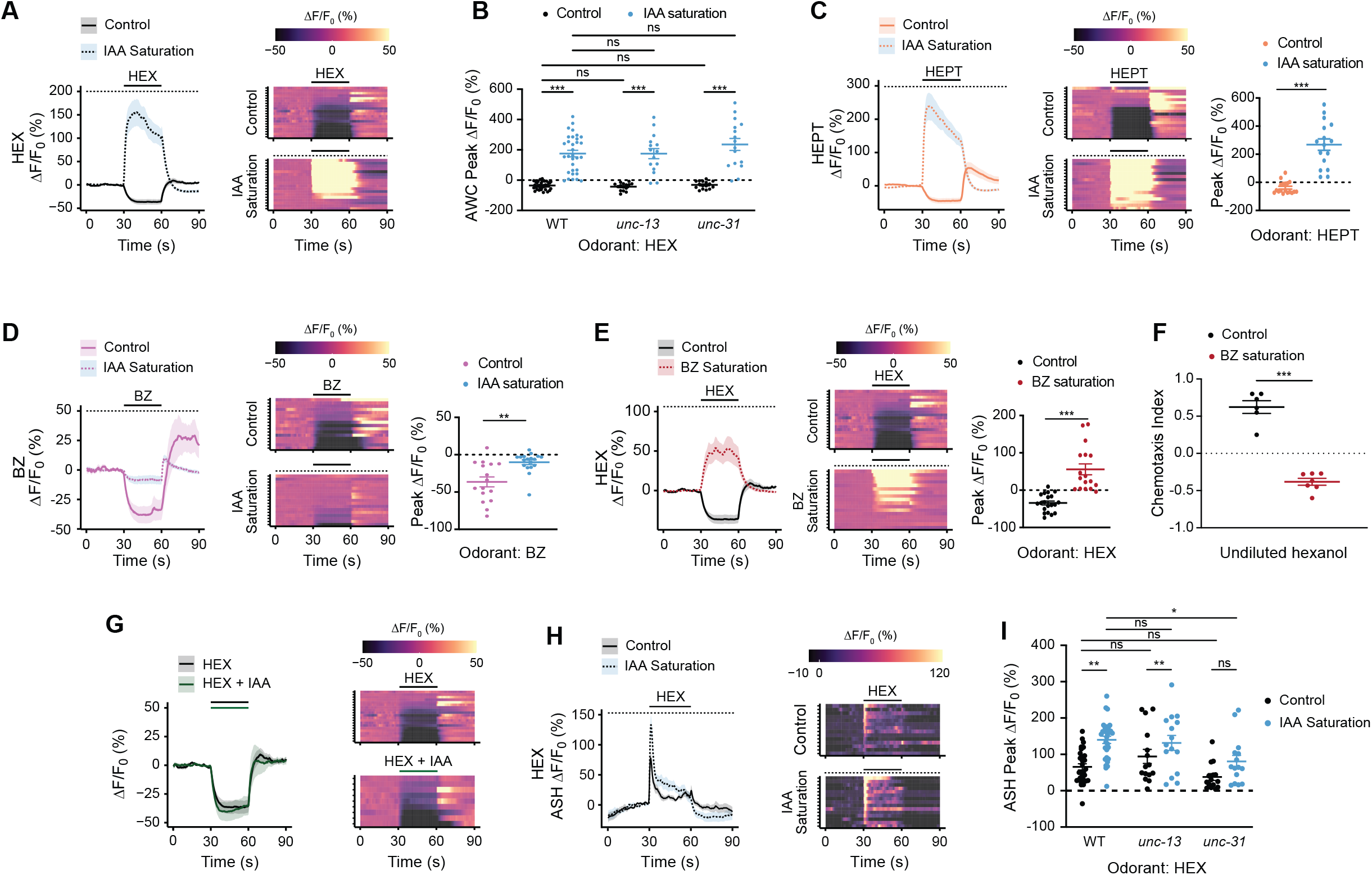
Hexanol-mediated inhibition of AWC is inverted to activation in saturating odor conditions. **A,C,D,E,G,H)** (Left) Average changes in GCaMP3 fluorescence in AWC (**A,C,D,E,G**) and ASH (**H**) in response to a 30s pulse of 10^−4^ dilution of the indicated odorant(s) (solid line). In **G**, odorants used were a mixture of a 10^−4^ dilution each of hexanol and IAA. The presence of saturating chemicals in the imaging chip at 10^−4^ dilution is indicated by a dashed line. Shaded regions are SEM. (Right in **A,G**, Center in **C,D,E**) Corresponding heatmaps of changes in fluorescence intensity. Each row in the heatmaps shows responses from a single AWC or ASH neuron from different animals; n≥15 each. (Right in **C,D,E**) Quantification of peak fluorescence intensity changes upon odorant onset under non-saturated or saturated conditions. Each dot is the response from a single neuron. Wild-type hexanol response data in control conditions were interleaved with experimental data in **A, E**, and **G** and are repeated. **B, I**) Quantification of peak fluorescence intensity changes upon odorant onset under non-saturated or saturated conditions in AWC (**B**) and ASH (**I**). Each dot is the response from a single neuron. Wild-type data are from **A** (for **B**) and **H** (for **I**). Average traces and heatmaps of responses in *unc-13(e51)* and *unc-31(e928)* mutants in AWC and ASH are shown in Figures S2C and S3B-E, respectively. **F)** Behavioral responses of animals to a point source of undiluted hexanol on plates with or without saturating benzaldehyde at 10^−4^ dilution. Each dot is the chemotaxis index of a single assay plate containing ~100-200 adult hermaphrodites. Assays were performed in duplicate over at least 3 days. Long horizontal bars indicate the mean; errors are SEM. *, **, ***: *P* <0.05, 0.01 and 0.001 and, respectively (**C-F**: Mann Whitney Wilcoxon test; **B,I**: Kruskal-Wallis with posthoc pairwise Wilcoxon test and Benjamini-Hochberg method for *P*-value correction); ns – not significant. HEX: 1-hexanol, HEPT: 1-heptanol, BZ: benzaldehyde.

We tested whether sIAA-dependent inversion of the response to medium-chain alcohols extends to additional AWC-sensed odorants. However, in sIAA, benzaldehyde decreased calcium levels in AWC albeit with a significantly reduced amplitude than in control conditions (Figure 2D). These responses are consistent with decreased attraction, but not aversion, to a point source of benzaldehyde in sIAA (Figure 1A). Similarly, upon saturation with hexanol, IAA elicited variable responses of smaller amplitude, but did not invert the response (Figure S2D), consistent with animals being indifferent to IAA in hexanol saturation conditions (Figure S1A). Calcium levels in AWC were also robustly decreased by the attractive chemical 2-methylpyrazine (Figure S2E). In sIAA, this chemical evoked variable responses of small amplitudes, but again did not increase intracellular calcium levels in AWC (Figure S2E). Attraction to this chemical was unaltered in sIAA (Figure S2F), likely due to this behavior being driven primarily by the AWA olfactory neuron pair (Bargmann et al., 1993). These results indicate that upon saturation, the observed response inversion upon odorant addition to AWC may be elicited only by specific chemicals such as hexanol and heptanol.

We also tested whether saturation by an AWC-sensed odorant other than IAA also results in hexanol-mediated activation of AWC. Indeed, we found that saturation with benzaldehyde resulted in hexanol-driven increases in calcium in AWC together with avoidance of hexanol (Figure 2E,F). In contrast, neither a hexanol/IAA mixture, nor pre-exposure to sIAA followed by a hexanol pulse, was sufficient to elicit an increase in calcium levels in AWC (Figure 2G, Figure S2G). We conclude that while attractive chemicals typically decrease intracellular calcium in AWC, upon saturation with an AWC-sensed attractant, hexanol and heptanol instead increase intracellular calcium levels in AWC. Activation of AWC correlates with avoidance of these medium-chain alcohols.

### ASH responds similarly to hexanol in the absence or presence of a saturating odor

Since our behavioral experiments indicate that the ASH neurons also contribute to hexanol avoidance (Figure 1D,E), we also examined hexanol-evoked responses in ASH. We verified that the transgenic strain expressing GCaMP3 in ASH exhibited behavioral responses to hexanol similar to those of wild-type animals (Figure S3A). Hexanol elicited a robust phasic response in ASH neurons with a rapid and transient rise upon hexanol addition (Figure 2H) (Fukuto et al., 2004; Hilliard et al., 2004). Response dynamics were similar in sIAA, although the response peak as well as the response baseline in the presence of hexanol were consistently higher as compared to the responses in control conditions (Figure 2H,I).

Hexanol responses in ASH in *unc-13* synaptic transmission mutants were similar to those in wild-type animals (Figure 2I, Figure S3C). However, in *unc-31* mutants defective in neuropeptidergic signaling (Sieburth et al., 2007; Speese et al., 2007), only a subset of animals responded to hexanol in control conditions although a larger fraction responded in sIAA (Figure 2I, Figure S3D). Moreover, the dynamics of the hexanol response in sIAA were altered such that the response appeared to be tonic (Figure 2I, Figure S3D). We tested whether peptidergic signaling from AWC might modulate hexanol response dynamics in ASH. Due to technical limitations, we could not assess hexanol responses in ASH in AWC-ablated animals. Sensory responses in AWC (including hexanol responses, see Figure 4A) are abolished in animals mutant for the *tax-4* cyclic nucleotide-gated channel (Bargmann et al., 1993; Komatsu et al., 1996). Hexanol responses in ASH in *tax-4* mutants resembled those in *unc-31* mutants (Figure S3E), suggesting that AWC or another *tax-4*-expressing neuron likely influences hexanol response dynamics in ASH. Together, these results indicate that unlike our observations in AWC, the hexanol response in ASH is not inverted in sIAA. However, while ASH responds cell-autonomously to hexanol, these responses may be modulated by AWC.

### The ODR-3 Gα protein is necessary for the hexanol driven response inversion in AWC in odorant saturation conditions

To probe the molecular mechanisms underlying the context-mediated hexanol response inversion in AWC, we tested the behaviors and hexanol responses of mutants previously implicated in AWC sensory signal transduction. Binding of odorants to their cognate receptors in AWC decreases intracellular cGMP levels either via inhibiting the activity of receptor guanylyl cyclases such as ODR-1 and DAF-11, and/or by promoting the activity of one or more phosphodiesterases, via heterotrimeric G proteins (Figure 3A) (Ferkey et al., 2021). Reduced intracellular cGMP levels closes cGMP-gated channels, inhibits calcium influx, and promotes attraction (Figure 3A) (Chalasani et al., 2007; Ferkey et al., 2021). The inversion in the hexanol response in AWC in control and sIAA conditions indicates that hexanol likely acts via distinct molecular mechanisms in AWC under different conditions to evoke a response.

**Figure 3.**
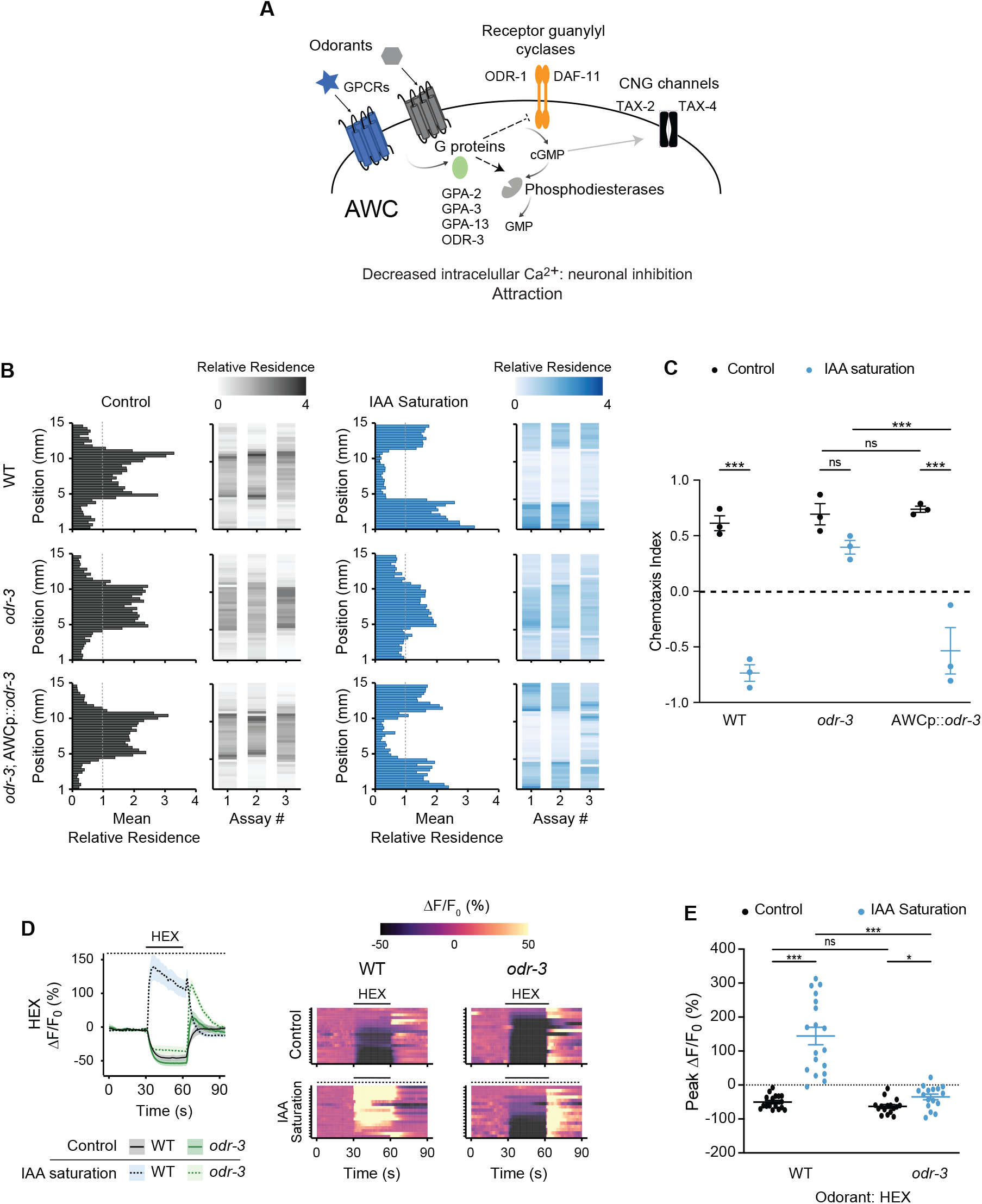
The ODR-3 Gα protein is required for hexanol-mediated activation of AWC in sIAA. **A)** Cartoon of the olfactory signal transduction pathway in AWC. **B)** Average residence histograms and heatmaps as described for Figure 1D. n=20-30 animals per assay; 3 biological replicates. An *odr-3* cDNA was expressed in AWC under the *odr-1* promoter. **C)** Chemotaxis indices calculated from behavioral assays shown in **B.** Each dot is the chemotaxis index from a single assay in behavior chips. **D)** (Left) Average changes in GCaMP3 fluorescence in AWC in response to a pulse of 10^−4^ hexanol in wild-type and *odr-3(n2150)* mutants. The presence of saturating chemicals in the imaging chip at 10^−4^ dilution is indicated by a dashed line. Shaded regions are SEM. (Right) Corresponding heatmaps of changes in fluorescence intensity; n≥15 each. Wild-type hexanol response data in sIAA were interleaved with experimental data in Figure 4C and are repeated. **E)** Quantification of fluorescence intensity changes upon hexanol odorant onset under nonsaturated or saturated conditions from data shown in **D**. Each dot is the response from a single neuron. Long horizontal bars indicate the mean; errors are SEM. *, ***: *P*<0.05 and 0.001, respectively (**C**: two-way ANOVA with Bonferroni’s correction; **E**: Kruskal-Wallis with posthoc pairwise Wilcoxon test and Benjamini-Hochberg method for *P*-value correction); ns – not significant.

While the hexanol and IAA receptors in AWC are unknown, multiple Gα proteins are expressed in AWC and have been implicated in mediating odorant signal transduction in this neuron type (Figure 3A) (Ferkey et al., 2021; Jansen et al., 1999; Lans et al., 2004; Roayaie et al., 1998). Loss of function mutations in the ODR-3 Gα_i_/Gα_o_-like protein decreases although does not fully abolish attraction to multiple AWC-sensed chemicals including IAA (Kato et al., 2014; L’Etoile and Bargmann, 2000; Lans et al., 2004; Roayaie et al., 1998). *odr-3* null mutants continued to be attracted to hexanol, indicating that this Gα protein is partly dispensable for this behavior (Figure 3B,C, Figure S4A). Animals mutant for the additional AWC-expressed nematode-specific Gα genes *gpa-2, gpa-3*, and *gpa-13*, as well as animals triply mutant for all three *gpa* genes, also retained the ability to be attracted to hexanol (Figure S4A). In contrast, *odr-3*, but not the *gpa* single or triple mutants, no longer avoided hexanol in sIAA, but were instead attracted, similar to the behaviors of these animals under control conditions (Figure 3B,C, Figure S4A). *gpa-3 gpa-13 odr-3* triple mutants were also attracted to hexanol in sIAA (Figure S4A). Hexanol avoidance behavior was rescued upon expression of *odr-3* specifically in AWC (Figure 3B,C).

To correlate neuronal responses with behavior, we next examined hexanol-evoked intracellular calcium dynamics in AWC in *odr-3* mutants. While hexanol decreased intracellular calcium concentrations in AWC similarly in both wild-type and *odr-3* animals, hexanol failed to increase calcium levels in AWC in sIAA in *odr-3* mutants (Figure 3D,E). Instead, hexanol continued to inhibit AWC in these animals (Figure 3D,E) consistent with *odr-3* mutants retaining attraction to hexanol in sIAA. A trivial explanation for the observed phenotype is that ODR-3 may be required for IAA-mediated saturation in AWC, leading to similar hexanol-evoked responses in *odr-3* mutants in both unsaturated and saturated conditions. Consistent with previous work showing that AWC retains partial ability to respond to IAA (Kato et al., 2014; Yoshida et al., 2012), IAA decreased intracellular calcium levels in AWC, although both the response amplitude as well as the number of responding neurons were reduced as compared to responses in wild-type animals (Figure S4B). We conclude that while hexanol attraction does not require ODR-3, ODR-3 is essential for the hexanol-mediated activation of AWC in sIAA. However, we are unable to exclude the possibility that IAA does not fully and/or bilaterally saturate AWC in *odr-3* mutants leading to the observed defect in hexanol-evoked activation of this neuron type.

### Hexanol-mediated inhibition but not activation of AWC requires the ODR-1 receptor guanylyl cyclase

We next asked whether hexanol acts via distinct downstream effector pathways in AWC in the absence or presence of sIAA to decrease or increase intracellular calcium levels, respectively. Both activation and inhibition by hexanol were abolished in animals mutant for the *tax-4* cyclic nucleotide-gated channel, indicating that both responses require cGMP signaling (Figure 4A). Consistent with a demonstrated requirement for the ODR-1 receptor guanylyl cyclase in mediating attraction to IAA (Bargmann et al., 1993; L’Etoile and Bargmann, 2000; Shidara et al., 2017), *odr-1* mutants also failed to be attracted to hexanol and were instead weakly repelled (Figure 4B). However, *odr-1* mutant animals robustly avoided hexanol in sIAA (Figure 4B), indicating that ODR-1 is dispensable for hexanol avoidance, but is necessary for attraction.

**Figure 4.**
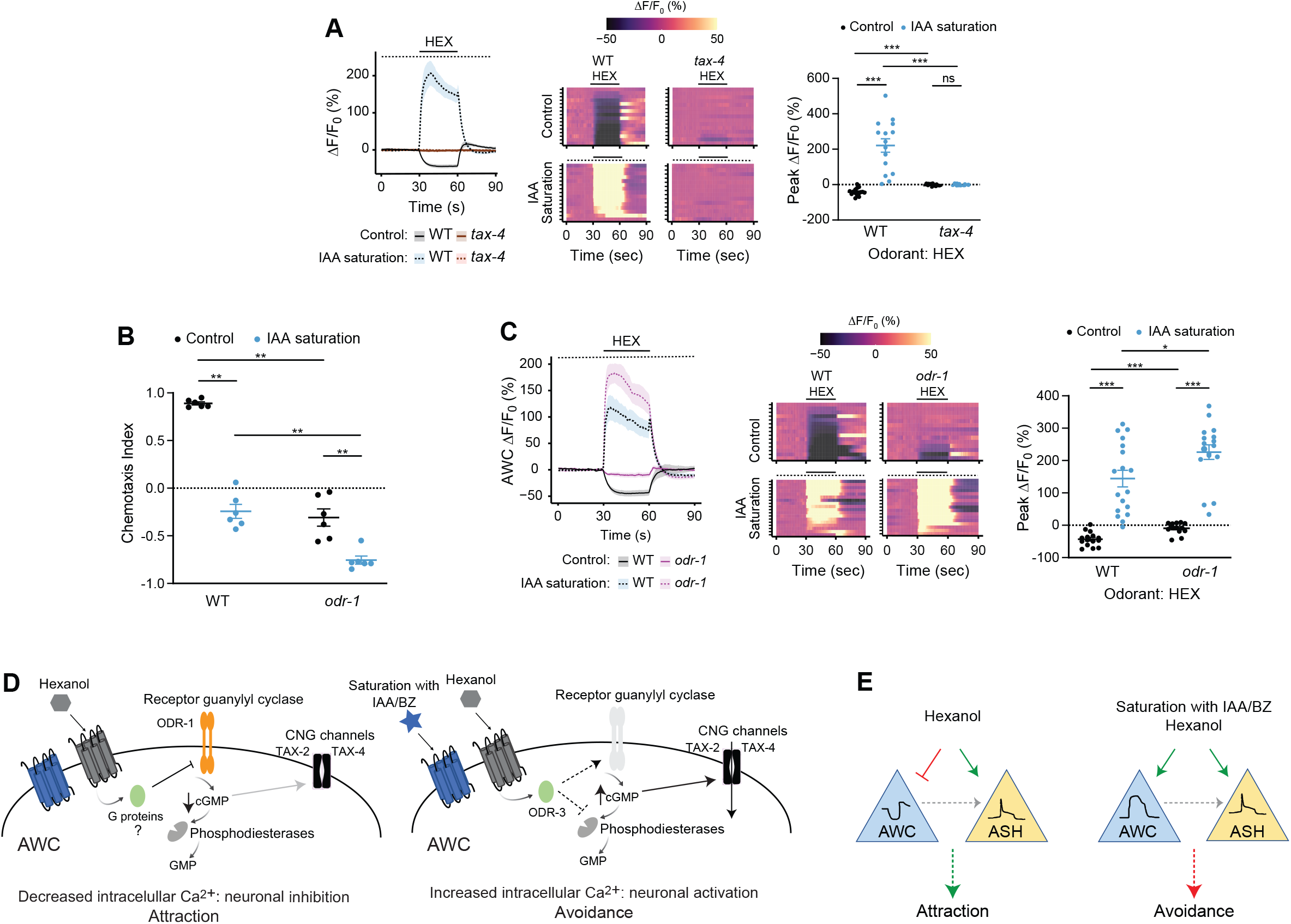
The ODR-1 receptor guanylyl cyclase is required for hexanol-evoked inhibition but not activation of AWC. **A,C)** (Left) Average changes in GCaMP3 fluorescence in AWC in response to a pulse of 10^−4^ hexanol in wild-type, *tax-4(p678)* and *odr-1(n1936)* animals. The presence of saturating chemicals in the imaging chip at 10^−4^ dilution is indicated by a dashed line. Shaded regions are SEM. (Center) Corresponding heatmaps of changes in fluorescence intensity are shown at right. Each row in the heatmaps shows responses from a single AWC neuron from different animals; n≥15 each. Control wild-type data were interleaved with experimental data in **A** and **C**; a subset of wild-type data in control conditions is repeated in these panels. (Right) Quantification of fluorescence intensity changes upon hexanol onset under non-saturated or saturated conditions. Each dot is the response from a single neuron. **B)** Behavioral responses of animals to a point source of 1:10 dilution of hexanol on plates with or without sIAA at 10^−4^ dilution. Each dot is the chemotaxis index of a single assay plate containing ~100-200 adult hermaphrodites. Assays were performed in duplicate over at least 3 days. Long horizontal bars indicate the mean; errors are SEM. *, **, ***: *P*<0.05, 0.01 and 0.001, respectively (Kruskal-Wallis with posthoc pairwise Wilcoxon test and Benjamini-Hochberg method for *P*-value correction). **D)** Proposed model for hexanol-mediated inhibition and activation of AWC in an odorant context-dependent manner. See text for details. **E)** In control conditions, hexanol inhibits AWC, and AWC-driven attraction predominates over ASH-driven avoidance. In saturation conditions, hexanol activates both AWC and ASH; either neuron can drive avoidance. AWC may modulate ASH hexanol responses via peptidergic signaling (gray dashed arrow).

Although ASH does not express *odr-1*, cGMP signaling in AWC has previously been shown to modulate ASH responses to a subset of nociceptive chemicals via a gap junction network (Krzyzanowski et al., 2016). In one model, hexanol responses could be lost in AWC in *odr-1* mutants regardless of odorant conditions, and hexanol avoidance could be driven by ASH alone. Alternatively, AWC may retain the ability to be activated but not inhibited by hexanol in *odr-1* mutants. To distinguish between these possibilities, we examined hexanol-evoked changes in calcium dynamics in AWC. Addition of either IAA or hexanol decreased intracellular calcium levels in AWC only to a minor extent in *odr-1* mutants (Figure 4C, Figure S4B), consistent with the inability of these animals to be attracted to either chemical. However, in sIAA, hexanol again robustly increased intracellular calcium levels in AWC in *odr-1* mutants (Figure 4C), correlated with these mutants retaining the ability to avoid hexanol. These results indicate that while hexanol acts via ODR-1 to inhibit AWC in control conditions and drive attraction, in sIAA, hexanol acts via an ODR-1-independent pathway to activate these neurons, and promote aversion.

## DISCUSSION

Here we show that the behavioral response of *C. elegans* to a food-related odor is inverted from attraction to avoidance in the continuous presence of a second attractive chemical. We find that this behavioral inversion is correlated with an inversion in the odorant response in a single olfactory neuron type via engagement of different intracellular signal transduction pathways in different chemical environments. Bidirectional responses of neurons such as parietal eye photoreceptors in lower vertebrates to blue or green light, and olfactory neurons of *Drosophila* in response to different odors have been reported (Cao et al., 2017; Hallem and Carlson, 2006; Hallem et al., 2004; Solessio and Engbretson, 1993; Su et al., 2006). In this work we describe a mechanism by which a single chemical evokes bidirectional sensory responses in a context-dependent manner in a single chemosensory neuron type, and suggest that related principles may underlie aspects of stimulus encoding and stimulus discrimination across sensory modalities.

In control conditions, we propose that hexanol acts via its cognate receptor(s) in AWC and Gα proteins other than or in addition to ODR-3 to inhibit the ODR-1 receptor guanylyl cyclase to close the TAX-2/TAX-4 channels and decrease intracellular calcium concentrations (Figure 4D) (Chalasani et al., 2007). However, in the presence of saturating AWC-sensed chemicals, the hexanol receptor instead likely acts primarily via the ODR-3 Gα protein to activate a receptor guanylyl cyclase other than ODR-1 (or inhibit a phosphodiesterase) to increase cGMP levels, open the TAX-2/TAX-4 channels and increase intracellular calcium to activate AWC (Figure 4D). The engagement of distinct signaling pathways in different odorant contexts suggests that the observed increase in hexanol/heptanol-evoked intracellular calcium levels is unlikely to simply be due to disinhibition, but instead represents a stimulus-driven neuronal response. Hexanol-evoked inhibition of AWC likely overrides the response in ASH to drive robust hexanol attraction in control conditions, but in saturating odor, activation of either AWC or ASH is sufficient to drive hexanol avoidance (Figure 4E). Branched and straight-chain alcohols are produced by multiple bacteria that are food sources for *C. elegans* (Elgaali et al., 2002; Worthy et al., 2018). The identities and temporal profiles of alcohols relative to other detected odorants may enable *C. elegans* to assess food quality and execute the appropriate behavioral decision.

How might hexanol engage different downstream effector pathways under different odorant conditions? Occupancy of a shared receptor by IAA may antagonize hexanol binding, and drive hexanol-mediated activation of a different signaling pathway via alternate AWC-expressed hexanol receptor(s). Antagonism of olfactory receptors by odorants in mixtures has now been extensively described and shown to play a role in stimulus encoding and odorant discrimination (Araneda et al., 2000; Oka et al., 2004; Pfister et al., 2020; Reddy et al., 2018; Xu et al., 2020; Zak et al., 2020). However, a mixture of IAA and hexanol does not activate AWC, and saturation with the structurally distinct chemical benzaldehyde is also sufficient to activate this neuron type. Although we are unable to exclude the possibility that AWC-sensed chemicals share a broadly tuned receptor that alters neuronal responses based on odorant context (MacWilliam et al., 2018; Turner and Ray, 2009), we favor the notion that saturation with an AWC-sensed chemical (and/or mutations in *odr-1*) alters neuronal state in a manner that then dictates the differential usage of intracellular signaling pathways by medium-chain alcohol receptor(s) to elicit distinct sensory responses. The as yet unidentified hexanol (and heptanol) receptor(s) may be modified in a neuronal state-dependent manner to promote coupling to distinct effector pathways upon ligand binding (Calebiro et al., 2021; Flock et al., 2017; Patwardhan et al., 2021). Alternatively, differential compartmentalization of signaling complexes within the AWC sensory cilia membrane may promote the usage of distinct signal transduction machinery in different neuronal conditions (Ellisdon and Halls, 2016; Hilgendorf et al., 2019; L’Etoile and Bargmann, 2000; Langeberg and Scott, 2015; Polit et al., 2020).

Attraction driven by AWC and other sensory neurons has previously been shown to be markedly reduced or switched to avoidance in specific mutant backgrounds or upon association with starvation and odorant experience (Adachi et al., 2010; Cho et al., 2016; Colbert and Bargmann, 1995; L’Etoile et al., 2002; Ohno et al., 2014; Tomioka et al., 2006; Tsunozaki et al., 2008). However, these forms of behavioral plasticity are regulated by modulation of synaptic transmission in the circuit, with little to no change in the primary sensory response. A potential advantage of differential usage of intracellular signaling pathways over modulation of sensory neuron synaptic output is the ability to discriminate between, and differentially respond to, each stimulus sensed by that neuron in a context-dependent manner. This mechanism is particularly advantageous for polymodal sensory neurons such as those in *C. elegans* (Ferkey et al., 2021) in which state-dependent engagement of different signaling pathways within a single sensory neuron type may allow animals to more effectively assess the salience of individual olfactory cues. Complex chemical response strategies have also been described in *Drosophila* gustatory neurons that express multiple ligand-gated ion channel receptors for different chemicals, and may represent a general mechanism by which organisms efficiently encode stimulus properties (Devineni et al., 2021; Stanley et al., 2021).

Chemical encoding strategies in *C. elegans* have largely been studied in response to monomolecular odorants. It will be important to expand this analysis to different odorant contexts and mixtures presented in different temporal sequences to more closely resemble environments that worms may encounter in the wild. As the signaling content of sensory cells across different organisms is described more fully (Kozma et al., 2020; McLaughlin et al., 2021; Taylor et al., 2021; van Giesen et al., 2020; Zheng et al., 2019), a challenging next step will be to assess how different intra- and intercellular pathways are used under different conditions, and how this response flexibility is translated through the circuit to drive adaptive behavioral responses.

## MATERIALS and METHODS

### Strains and growth conditions

All *C.elegans* strains were maintained on nematode growth medium (NGM) at 20°C. 5 days prior to behavioral assays, 10 L4 larvae per genotype were picked to 10 cm assay growth plates (day 1), and young adults were tested in behavioral and calcium imaging assays 4 days later (day 5). Animals were maintained under well-fed conditions at all times.

To standardize growth conditions, NGM plates were seeded with bacteria as follows: concentrated *Escherichia coli* OP50 was cultured by inoculating 10 μl of a starter OP50 culture (grown in LB for ~2 hr from a single colony) per 1L of SuperBroth media (3.2% w/v tryptone, 2.0% yeast extract, 0.5% NaCl). SuperBroth cultures grown overnight were treated with a low concentration of the antibiotic gentamicin (300 ng/ml; Sigma G1397) for ~4 hours, centrifuged for 20 minutes at 4°C, and the resulting pellets resuspended in 75 ml of S-Basal buffer. The concentrated bacterial food was stored at −80°C and thawed as needed to seed plates (1 ml/10cm plate).

All strains were constructed using standard genetic procedures. The presence of mutations was confirmed by PCR-based amplification and/or sequencing. The *odr-1*p*::odr-3::SL2::mCherry* (PSAB1269) plasmid was injected at 10 ng/μl together with the *unc-122*p*::gfp* co-injection marker at 50 ng/μl to generate transgenic rescue strains. Expression patterns and phenotypes were confirmed in initial experiments using multiple independent transgenic lines, and a single line was selected for additional analysis. A complete list of strains used in this work is provided in Table S1.

### Molecular biology

An *odr-3* cDNA (Roayaie et al., 1998) was cloned into a worm expression plasmid containing ~1.0 kb upstream *odr-1* regulatory sequences using standard restriction enzyme cloning (PSAB1269).

### Plate chemotaxis assays

Chemotaxis assays were performed according to previously published protocols (Bargmann et al., 1993; Troemel et al., 1997). Assays were performed on 10 cm square or round plates with two 1 μl spots of odorant and the diluent ethanol at either end, together with 1μl of 1M sodium azide at each spot to immobilize worms. Odorants were diluted freshly in ethanol as needed. Saturation assays were performed using the same protocol, except that the relevant odorant was added to the assay agar before pouring plates (1 μl odorant at 10^−4^ dilution /10ml agar). Animals were washed off growth plates with S-Basal and washed twice subsequently with S-Basal and once with Milli-Q water. Washed animals were placed at the center of the assay plate and allowed to move for an hour. The number of worms in two horizontal rows adjacent to the odor and ethanol spots was quantified at the end of the assay. Each assay was performed at least in duplicate each day; data are reported from biologically independent assays performed on at least 3 days.

### Osmotic avoidance assay

Osmotic avoidance behavior assays were performed essentially as previously described (Cornils et al., 2016; Culotti and Russell, 1978). 10 young adult worms were transferred without food to an agar plate and allowed to recover for at least 2 min. They were then placed in the center of an NGM plate with a ring of 8M glycerol containing bromophenol blue (Sigma B0126). The number of worms inside and outside of the ring was counted after 10 minutes.

### Microfluidics behavioral assays

Microfluidics assays were performed following published protocols, using custom designed microfluidic devices (https://github.com/SenguptaLab/HEXplasticity.git) (Albrecht and Bargmann, 2011). The assembled microfluidic device was degassed in a vacuum desiccator for ~30 minutes prior to loading a 5% w/v poloxamer surfactant (Sigma P5556) with 2% xylene cyanol (2 mg/ml) solution through the outlet port. These steps ensured that the arena was bubble-free prior to loading the worms and stimulus reservoirs. Buffer and stimulus flowed by gravity from elevated reservoirs and were controlled with manual Luer valves. 20-30 young adult animals were transferred to unseeded plates and flooded with S-Basal buffer to remove any residual bacteria. The worms were then transferred into a tube and gently loaded into the buffer-filled arena via syringe. After allowing the worms to disperse throughout the arena (~5 min), the flow of the odorant stimulus was started. 3 parallel stripes flowed through the stimulus; 2 outer stripes consisted of buffer and the central stripe contained the odorant. 2% xylene cyanol (2 mg/ml) was added to the odorant to allow visualization and tracking. For IAA saturation assays, in addition to the stimulus odorant in the middle stripe, 10^−4^ IAA was included in all buffer and stimulus reservoirs. Movies were recorded at 2 Hz on a PixelLink camera while worms were exposed to 20 minutes of constant odor. Following each experiment, the devices were flushed with water and soaked in ethanol overnight to remove any residual odorant. Prior to using the devices for additional assays, the chip was rinsed in water and baked at 50°C for a minimum of 4 hours to evaporate any residual ethanol odor and liquid. The cleaning procedure was validated by buffer-buffer control assays, in which worms showed no spatial preference.

All movie acquisition, processing, and subsequent behavioral analysis was performed via custom MATLAB software modified from (Albrecht and Bargmann, 2011) (https://github.com/SenguptaLab/HEXplasticity.git). Data visualization and figures were generated using RStudio (version 1.3.959). A minimum of 3 assays per condition were performed on multiple days, and mean relative residency and chemotaxis index in respect to spatial stimuli was calculated. Briefly, the *y*-position data were binned into 50 bins for each assay. Relative residence was calculated by counting the # of tracks in each of the 50 *y*-position bins, and dividing each of these counts by the average # of counts across all 50 bins. Average residency histograms show the average of these residency values for each *y*-position bin over 3 assays/condition. Chemotaxis index was calculated as (normalized # of tracks within the odorant – normalized # of tracks in buffer)/total # of normalized tracks. Track numbers were normalized by calculating the # tracks X (total length of arena/ length of respective buffer or odorant region). Stripe boundaries containing the odorant were determined using luminance data using xylene cyanol dye. To account for variable luminance across the device, luminance values were normalized using a linear regression fit. Boundaries were identified as the first and second sign switch of luminance values using the normalized luminance data.

### Calcium imaging

Calcium imaging was performed as previously described, using custom microfluidic devices (Chronis et al., 2007; Neal et al., 2015). Imaging was conducted on an Olympus BX52WI microscope with a 40X oil objective and Hamamatsu Orca CCD camera. Recordings were performed at 4 Hz. All odorants were diluted in S-Basal buffer and 1 μl of 20 μM fluorescein was added to one of the channels to confirm correct fluid flow. IAA saturation assays included IAA (1:10,000) in all channels, including the worm loading buffer. 1 mM (-)- tetramisole hydrochloride (Sigma L9756) was added to the S-Basal buffer to paralyze body wall muscles and keep animals stationary. To prevent the chip from clogging, poloxamer surfactant (Sigma P5556) was also added to S-Basal while loading the worms. Odor evoked calcium transients in AWC and ASH sensory neurons were similar in the presence or absence of these chemicals. AWC and ASH neurons were imaged for one cycle of 30s buffer/30s odor/30s buffer stimulus. For pre-exposure experiments, animals were transferred to either control or IAA-saturated chemotaxis plates. After 1 hour on the chemotaxis plates, animals were loaded in the calcium imaging chip. Each animal was imaged within ~5 minutes after removal from the plate. Imaging was also performed in the presence of buffer only to ensure that observed neuronal responses were to the odor stimulus and not artifactual.

Recorded image stacks were aligned with Fiji using the Template Matching plugin, and cropped to a region containing the cell body. The region of interest (ROI) was defined by outlining the desired cell body; background subtracted fluorescence intensity of the ROI was used for subsequent analysis. To correct for photobleaching, an exponential decay was fit to fluorescence intensity values for the first 30s and the last 20s of imaging (prior and post stimulus). The resulting curve was subtracted from original intensity values. Amplitude was calculated as maximum change in fluorescence (F-F_0_) in the 10s following odor addition; F_0_ was set to the average ΔF/F_0_ value for 5s before odor onset. Data visualization and figures were generated using RStudio (version 1.3.959). Photomask designs for customized microfluidic olfactory chips adapted from (Chronis et al., 2007) are available on GitHub (https://github.com/SenguptaLab/HEXplasticity.git). Reported data were collected from biologically independent experiments over at least 2 days.

### Statistical analyses

Excel (Microsoft) and GraphPad Prism version 9.0.2 (www.graphpadpad.com) were used to generate all chemotaxis plate assay data. Plate chemotaxis index and peak ΔF/F_0_ amplitude data were analyzed using the Mann Whitney Wilcoxon or Kruskal-Wallis tests followed by the posthoc pairwise Wilcoxon test and Benjamini-Hochberg method for *P*-value correction. Chemotaxis index data derived from microfluidics behavioral assays were analyzed using oneway or two-way ANOVA followed by Bonferroni’s multiple comparison test. All statistical analyses were performed in R (https://www.R-project.org/) and RStudio (http://www.rstudio.com) and GraphPad Prism version 9.0.2 (www.graphpadpad.com)’. The tests used are indicated in the corresponding Figure Legends.

## ACKNOWLEDGEMENTS

We are grateful to the *Caenorhabditis* Genetics Center and Steven Flavell, Yun Zhang, and Gert Jansen for providing strains, the Brandeis Materials Research Science and Engineering Center (MRSEC) for access to the microfabrication facility, and Dirk Albrecht and Till Hartmann for assistance with analyses of microfluidics behavioral assays. We thank members of the Sengupta lab and Oliver Hobert for critical comments on the manuscript. This work was supported in part by the NSF (IOS 165518 and IOS 20142100 – P.S., MRSEC 2011846 to Brandeis University), and the NIH (F31 DC015186 - A.H.H.).

## AUTHOR CONTRIBUTIONS

Conceptualization: A.H.H., N.D.D., C.I.B., P.S.; Methodology: M.K., A.H.H., M.P.O’D.; Investigation: M.K., M.P.; Software: A.H.H., M.P.O’D.; Writing – original draft: M.K., P.S.; Writing – review and editing: M.K., A.H.H., M.P.O’D., M.P., N.D.D., C.I.B., P.S.; Visualization: M.K., A.H.H.; Supervision: P.S.; Funding acquisition: P.S.

## SUPPLEMENTAL FIGURE LEGENDS

**Figure S1.**
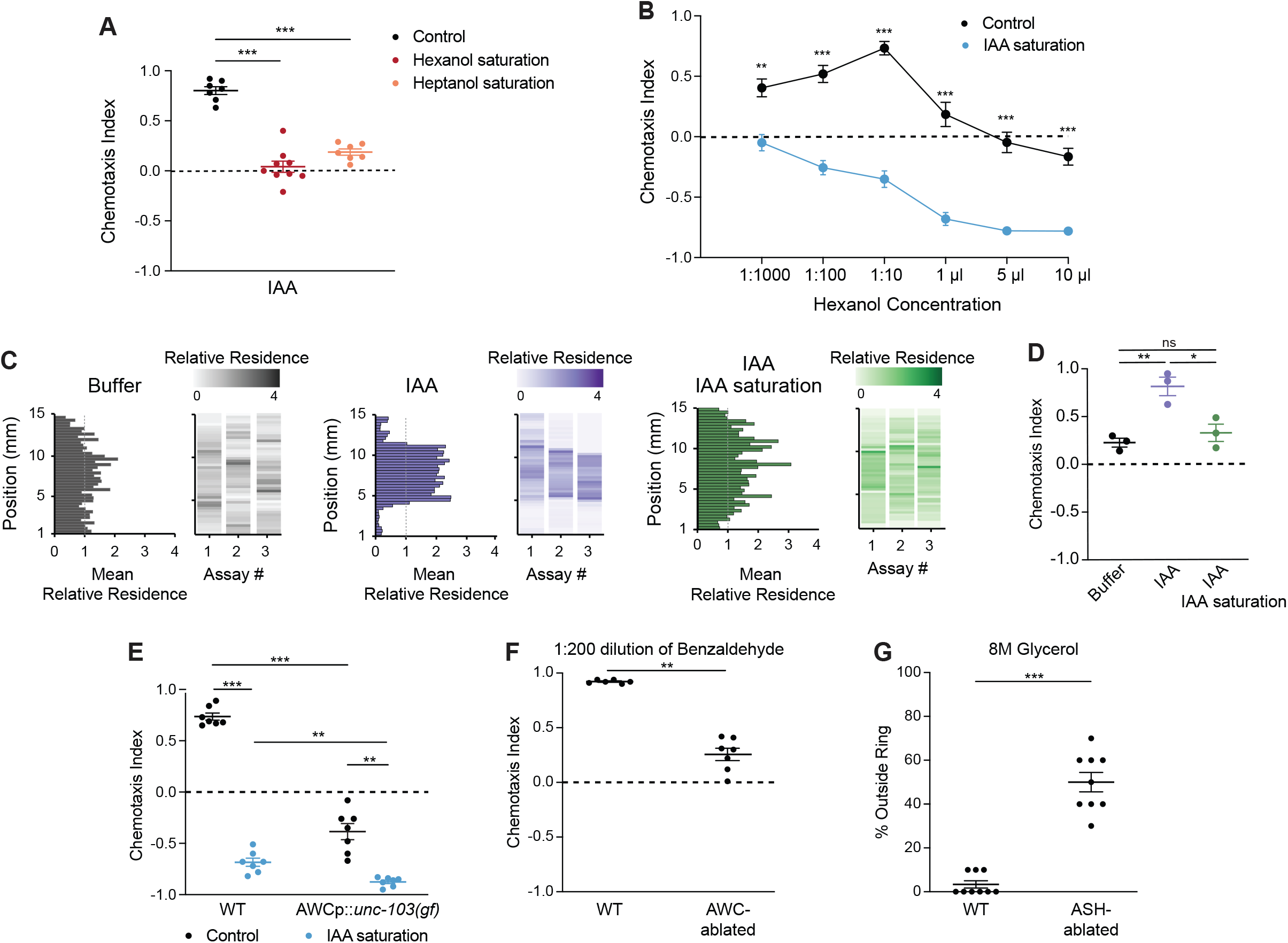
Attraction to hexanol is switched to aversion in sIAA. **A)** Behaviors of wild-type animals on control plates or plates saturated with either hexanol or 1-heptanol. Test odorant: 1:1000 dilution of IAA. **B)** Behaviors of wild-type animals on control or IAA-saturated plates to the indicated concentrations of hexanol. **C)** Average residence histograms and heatmaps as described for Figure 1D. n=20-30 animals per assay; 3 biological replicates. **D)** Chemotaxis indices calculated from behavioral assays shown in **C**. **E)** Behaviors of animals of the indicated genotypes on control or sIAA plates to undiluted hexanol. The *odr-1* promoter was used to drive *unc-103(gf)* in AWC (Yeon et al., 2021). **F)** Behaviors of animals of the indicated genotypes to a 1:200 dilution of benzaldehyde. **G)** Shown is the percentage of animals of the indicated genotypes that escaped a ring of 8M glycerol. In **A,E,F**, each dot represents the chemotaxis index of a single assay plate containing ~100-200 adult hermaphrodites. Assays were performed in duplicate on at least 3 days. In **B**, each dot represents the average chemotaxis index of 3-4 independent assays of ~100-200 animals each. In **D**, each dot represents the chemotaxis index from a single assay in behavior chips. In **G**, each dot represents a single osmotic avoidance assay of 10 animals. Long horizontal bars indicate the mean; errors are SEM. Errors are SEM. *, **, ***: *P*<0.05, 0.01 and *P*<0.001, respectively (**A,B,E**, Kruskal-Wallis with posthoc pairwise Wilcoxon test and Benjamini-Hochberg method for *P*-value correction, **F,G**, Mann Whitney Wilcoxon test, **D**, two-way ANOVA with Bonferroni’s correction).

**Figure S2.**
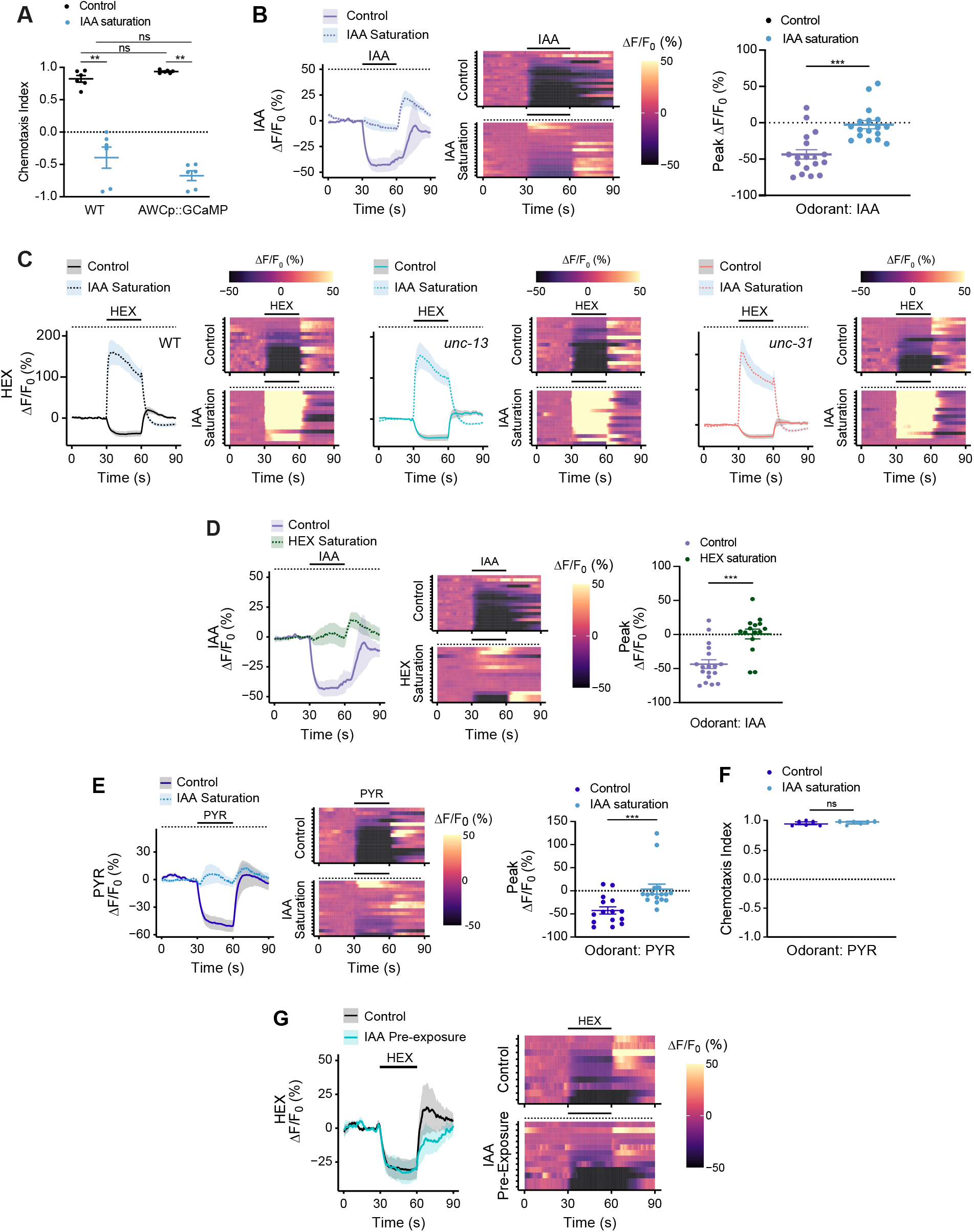
Hexanol but not other odorants activate AWC under odorant saturation conditions. **A,F)** Behaviors of wild-type animals or animals expression GCaMP3 under the *odr-1* promoter in AWC to a point source of 1:10 dilution of hexanol (**A**) or 1 μl of 10 mg/ml dilution of 2-methylpyrazine (PYR) (**F)** on plates with or without sIAA at 10^−4^ dilution. Each dot is the chemotaxis index of a single assay plate containing ~100-200 adult hermaphrodites; assays were performed in duplicate over at least 3 days. Long horizontal bars indicate the mean; errors are SEM. **: *P*<0.01 (**A**, Kruskal-Wallis with posthoc pairwise Wilcoxon test and Benjamini-Hochberg method for *P*-value correction, **F**, Mann Whitney Wilcoxon test); ns – not significant. **B-E,G)** (Left) Average changes in GCaMP3 fluorescence in AWC in response to a pulse of 10^−4^ dilution of the indicated odorants (solid line). The presence of saturating chemicals in the imaging chip at 10^−4^ dilution is indicated by a dashed line. In **G**, animals were pre-exposed to a 10^−4^ dilution of IAA (see Methods). Shaded regions are SEM. (Right in **C,G**, Center in **B,D,E**) Corresponding heatmaps of changes in fluorescence intensity. Each row in the heatmaps shows responses from a single AWC neuron from different animals; n≥14 each. (Right in **B,D,E**) Quantification of peak fluorescence intensity changes upon odorant onset under non-saturated or saturated conditions. Each dot represents the response from a single neuron. Control wild-type IAA response data were interleaved with experimental data in **B** and **D** and are repeated. Long horizontal bars indicate the mean; errors are SEM. ***: *P*< 0.001, (Mann Whitney Wilcoxon test). Quantification of responses in **C** are shown in Figure 3B. PYR – 2-methyl-1-pyrazine. Alleles in **C** were *unc-13(e51)* and *unc-31(e928)*.

**Figure S3.**
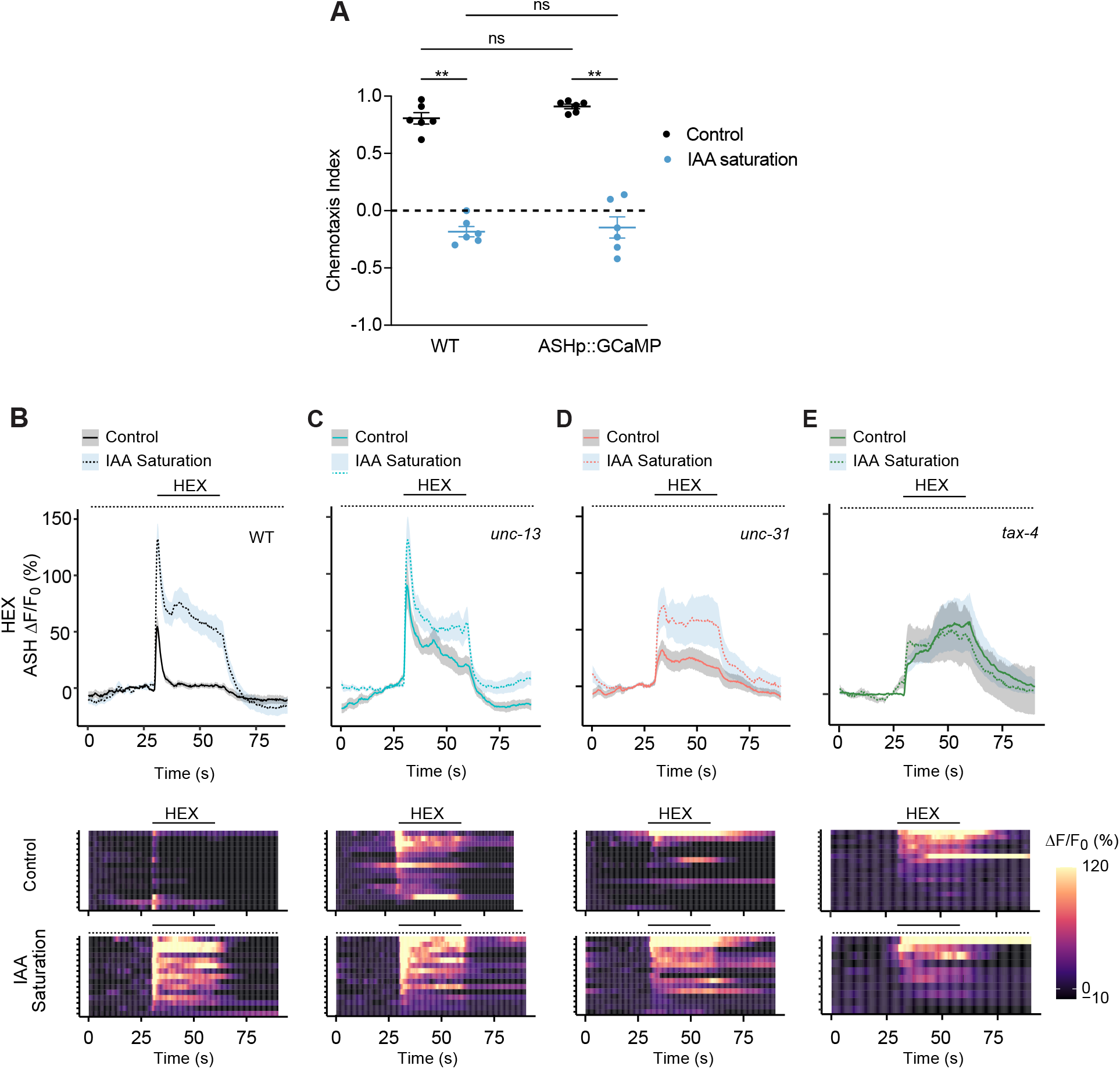
ASH responds partly cell-autonomously to hexanol. **A)** Behaviors of wild-type or a strain expressing GCaMP3 in ASH under the *sra-6* promoter to a point source of undiluted hexanol on plates without or with sIAA at 10^−4^ dilution. Each dot represents the chemotaxis index of a single assay plate containing ~100-200 adult hermaphrodites; assays were performed in duplicate over at least 3 days. Long horizontal bars indicate the mean; errors are SEM. **: *P*<0.01(Kruskal-Wallis with posthoc pairwise Wilcoxon test and Benjamini-Hochberg method for *P*-value correction); ns – not significant. **B-E)** (Top) Average changes in GCaMP3 fluorescence in ASH in response to a pulse of 10^−4^ dilution of hexanol (solid line) in animals of the indicated genetic backgrounds. The presence of saturating chemicals in the imaging chip at 10^−4^ dilution is indicated by a dashed line. Shaded regions are SEM. (Bottom) Corresponding heatmaps of changes in fluorescence intensity. Each row in the heatmaps shows responses from a single ASH neuron from different animals. n = 10 (*tax-4*), 15 (all other genotypes). Alleles used were *unc-13(e51), unc-31(e928)*, and *tax-4(p678)*. Quantification of peak response amplitudes are shown in Figure 2I.

**Figure S4.**
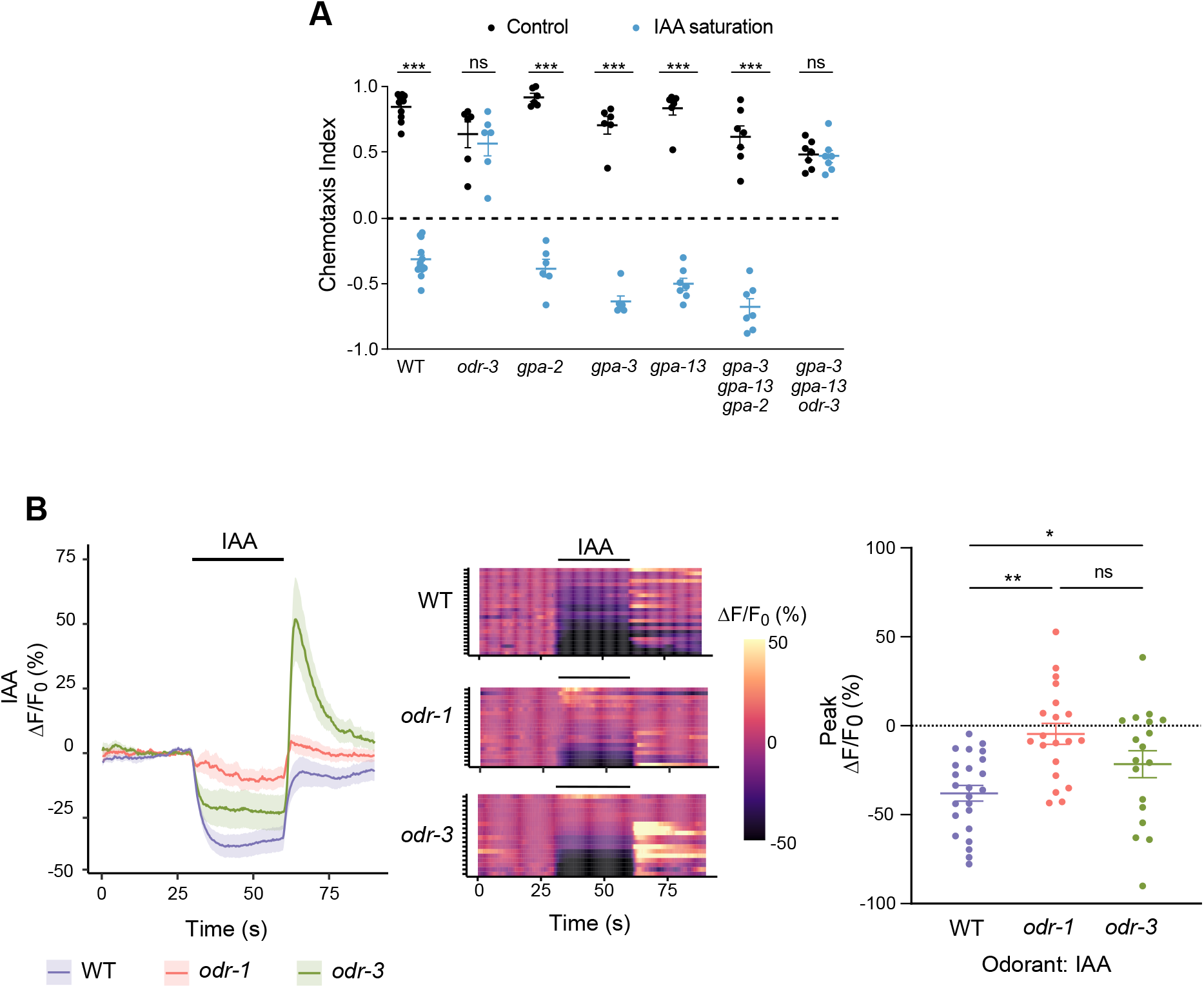
Hexanol acts via distinct signaling pathways to inhibit or activate AWC in a contextdependent manner. **A)** Behaviors of animals of the indicated genotypes (see Table S1) to a point source of 1:10 dilution of hexanol on plates with or without sIAA at 10^−4^ dilution. Each dot is the chemotaxis index of a single assay plate containing ~100-200 adult hermaphrodites. Assays were performed in duplicate over at least 3 days. Long horizontal bars indicate the mean; errors are SEM. ***: *P*<0.001 (Kruskal-Wallis with posthoc pairwise Wilcoxon test and Benjamini-Hochberg method for *P*-value correction); ns – not significant. **B)** (Left) Average changes in GCaMP3 fluorescence in AWC in response to a 30s pulse of 10^−4^ dilution of IAA (solid line) in animals of the indicated genetic backgrounds. The presence of saturating chemicals in the imaging chip at 10^−4^ dilution is indicated by a dashed line. Alleles used were *odr-1(n1936)* and *odr-3(n2150)*. Shaded regions are SEM. (Center) Corresponding heatmaps of changes in fluorescence intensity. Each row in the heatmaps shows responses from a single AWC neuron from different animals; n≥15 each. (Right) Quantification of peak response amplitudes. Each dot is the response from a single neuron. Long horizontal bars indicate the mean; errors are SEM. *, **: *P*<0.05 and 0.001, respectively (Kruskal-Wallis with posthoc pairwise Wilcoxon test and Benjamini-Hochberg method for *P*-value correction); ns – not significant.

**Video S1.** Wild-type animals are attracted to, or avoid, hexanol in the absence or presence of sIAA in microfluidics behavioral arenas. Video shows the locomotor responses of wild-type worms to odor stripes of 10^−4^ hexanol (HEX) in control (left) and sIAA (HEX/sIAA, right) conditions in a microfluidics behavioral chip. Odor stripes contain dye for visualization. Video is accelerated 40X.

**Video S2.** Hexanol decreases or increases intracellular calcium in AWC in the absence or presence of sIAA, respectively. A wild-type animal expressing the *odr-1*p*::GCaMP3* transgene was subjected to a 30s pulse of 10^−4^ hexanol (HEX onset) in control (HEX, left) and sIAA (HEX/sIAA, right) conditions. Video is accelerated 6X.

**Table S1.**
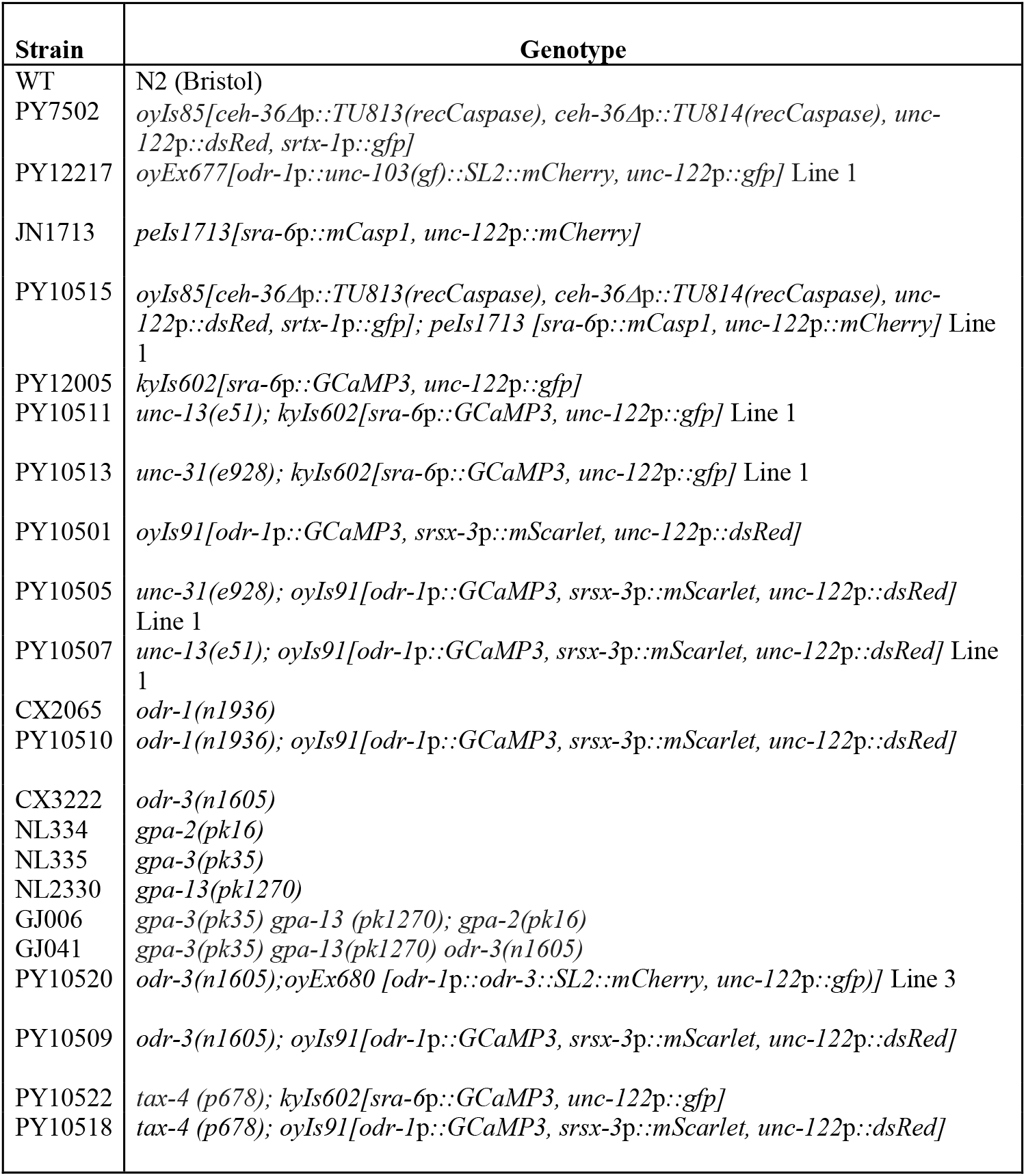
Strains used in this work.

